# Investigation of oxidative processes and revealing the changes in purine nucleoside phosphorylase activity levels in model animals in the development of diabetes mellitus

**DOI:** 10.1101/2022.01.26.477853

**Authors:** G.S Ghazaryan, L.M. Hovsepyan, L.G. Poghosyan, H. V. Zanginyan

## Abstract

Diabetes mellitus is a global and one of the most common healthcare problems. The immune system, having a certain autonomy in the recognition and removal of foreign cells, antigens and other compounds is also under strict homeostatic control, in which a lot of/ variety of biochemical reactions take part. And since the problem of treatment of diabetes mellitus is still not solved, it requires detailed research of different elements of pathogenesis at the molecular level. Taking into account the fact that the connection/ link between immune processes and enzymes of purine metabolism is doubtless, it is of interest to investigate the changes in purine metabolism. In previous studies a frequent combination and interrelation of purine and carbohydrate metabolism was shown. Aim of this study was the investigation of the enzymatic activity of PNP in serum, liver and in pancreas, and also determination of levels of lipid peroxidation indicators, NO content in blood plasma, glutathione peroxidase activity in the mitochondrial fraction of the pancreatic tissue of white rats with modeled alloxan diabetes. The obtained results have demonstrated statistically significant changes in the PNP activity level indicators in the investigated organs and stages of the disease. The indicators of oxidative processes MDA, NO and changes of the activity of GPx and PNP can be used as additional paraclinical indicators for early diagnostics and for detection and refinement of the level of activity of the pathologic process, its development and stage of the disease, which contributes to the appointment of individualized adequate therapy.

## INTRODUCTION

Diabetes mellitus is a global and one of the most common healthcare problems. The immune system, having a certain autonomy in the recognition and removal of foreign cells, antigens and other compounds is also under strict homeostatic control, in which a lot of/ variety of biochemical reactions take part. [1] And since the problem of treatment of diabetes mellitus is still not solved, it requires detailed research of different elements of pathogenesis at the molecular level. Taking into account the fact that the connection/ link between immune processes and enzymes of purine metabolism is doubtless, it is of interest to investigate the changes in purine metabolism[2]. In previous studies a frequent combination and interrelation of purine and carbohydrate metabolism was shown [3].

PNP is one of the most important enzymes of purine metabolism, characterizing the immune status of the organism. Inhibition of this enzyme leads to disruption of nucleoside homeostasis, which causes T-cell immunodeficiency [4]. Besides, as shown in one of the large-scale joint international projects on the use of proteomic methods for the identification of markers of hepatotoxicity induced by various agents, PNP together with vitamin D-binding protein, malate dehydrogenase, paraoxonase, cellular retinol-binding protein and F-protein is identified among the six earliest and most effective serum markers of hepatotoxicity [5]. In diabetes mellitus there are conditions for formation of oxidative stress: amount of oxidative substrates increases (glucose and lipids), the formation and activity of natural antioxidant systems - such as glutathione, superoxide dismutase, catalase and glutathione peroxidase decreases [10,11].

Aim of this study was the investigation of the enzymatic activity of PNP in serum, liver and in pancreas, and also determination of levels of lipid peroxidation indicators, NO content in blood plasma, glutathione peroxidase activity in the mitochondrial fraction of the pancreatic tissue of white rats with modeled alloxan diabetes.

## METHODS

The research was conducted on white <u>outbread</u> rats of 170-200 g body mass, contained in accordance with the rules of the European Convention for the Protection of Vertebrates Used for Experimental and Scientific Purposes (Strasbourg, 1986).

The rats were kept in controlled conditions providing 12-hour cycles of light and darkness in a temperature-controlled animal room (22 ° C). The rats had access to food and water. All rats received a standard laboratory diet (B&K Universal). The rats were divided into two groups: a control group of six animals and animals with induced alloxan diabetes. The amount of animals was chosen and calculated as n = 6 which corresponds to the minimal number of animals necessary for conducting adequate and good controlled investigation for performing precise and reliable statistical analysis of the obtained data.

The protocol on animals was approved by the Animal Ethics Committee of the Institute of Molecular Biology of the National Academy of Sciences of the Republic of Armenia and experiments were conducted in accordance with its guidelines and rules. All rats were anesthetized by intra-abdominal administration of the MMB mixture (0,3 mg / kg of body mass medetomidine, 4 mg/ kg midazolam, 5 mg/ kg butorfanol). Adequate depth of anesthesia was provided by testing of paw retraction and palpebral reflexes. All rats were killed by an overdose of carbon dioxide, followed by blood sampling from the carotid artery and subsequent decapitation.

Alloxan diabetes (AD) was caused by an injection of 40 mg/ kg of alloxan. The level of glucose in the capillary blood was detected with glucometer on 10, 15 and 20th days after alloxan injection. In the experiment, animals with sugar content more than 11-14 mmol/l were used.

After decapitation blood samples were obtained and organs - liver and pancreas were isolated for analysis. The blood after coagulation was centrifuged in a refrigerated centrifuge at 12.000 rotation per minute for 30 minutes and the obtained serum was used the same day for assessment of enzymatic activity. Pancreas and liver were washed from blood with chilled saline solution, homogenized in 0,1 M tris extraction solution pH 7,2, containing 5mM DTT and 1 mM EDTA and the obtained organ extracts were centrifuged at 18 thousand rpm for 30 minutes. Further the supernatants were used for assessment of enzymatic activity. PNP activity was assessed by guanine accumulation, which amount was assessed by color reaction with Folin reagent and was expressed in mmol of the substrate used per 1 g of wet tissue per minute, and for blood serum – mmol/l/min. Duration of the incubation in a water thermostat at 37 degree for assessment of the enzymatic activity was 6 minutes. The measurements were done spectrophotometrically, using quartz cuvettes at 750 nm wavelenghth [5].

Mitochondrial fraction was isolated in a medium containing 0,25 M saccharose and 0, 01M tris buffer, pH - 7,4, by a method of differential centrifugation on a K-24 centrifuge. Glutathione peroxidase activity was assessed by reduction reactions of hydrogen peroxide (H2O2) and lipid hydroperoxides (ROOH) in the presence of GSH [12, 15].

LPO activity was evaluated by the amount of Malondialdehyde (MDA) in the general homogenate and mitochondrial fraction of pancreas of control and alloxan diabetic rats. MDA was assessed by a reaction with Thiobarbituric acid [16].

The final product nitric oxide (NO) was assessed with Greys reactive (1 % Sulfanilamideб 0,1 % naphthylenediamine, 2,5 % phosphoric acid), the absorption of the solution was measured at wavelength of 546 nm. Sodium nitrite was used as a standard [17].

### Statistical analyses

For statistical data analysis on PNP SPSS (Statistical Package for Social Science) package was used. The nature of the distribution of the obtained data was determined by the Kolmogorov-Smirnov criterion. The comparative analysis was performed using a nonparametric Mann-Whitney test. The differences were statistically significant when *p* ≤ 0,05 or *p* ≤ 0,01. Correlational analysis was conducted using a nonparametric Spearman test. In the work refrigeratory centrifuge, spectrophotometer LKB Biochrom ULTROSPECII (Sweden), pH-meter PL-600 mrc (Israel), guanosine and ATP (Sigma).

## RESULTS AND DISCUSSION

Results of our investigation show that in case of alloxan-induced diabetes mellitus, there is a significant decrease in body weight of experimental animals and increase in daily water consumption (Table 2). Also, according to our observations the highest glucose level in blood was on day 20 (table 1).

**Table 1.** Dynamics of blood glucose level at experimental alloxan diabetes.

*Dynamics of blood glucose level at experimental alloxan diabetes*
| Experimental groups | Blood glucose level (mmol/L) |  |  |
| --- | --- | --- | --- |
|  | Days of experiment |  |  |
|  | 1 | 10 | 20 |
| Control | $3,8 \pm 0,2$ | $4,03 \pm 0,4$ | $3,9 \pm 0,03$ |
| Experimental group | $5,8 \pm 0,3$ | $8,3 \pm 0,4$ | $12,2 \pm 0,5$ |

**Table 2.** Dynamics of physiological indicators of laboratory animals at alloxan diabetes.

|  | Experimental groups |  |
| --- | --- | --- |
|  | Control group | experimental group |
| Body mass (g) | $250 \pm 10$ | $180 \pm 6$ |
| Water consumption (ml/day) | $20 \pm 3$ | $32 \pm 4$ |

Note: $p < 0,05$ — the reliability of the differences when comparing the indicators of the experimental groups with the control ones.

As the results of the research showed (Fig. 1), there are significant differences of the enzymatic activity between the investigated organs, liver and pancreas, however the similarity is also visible. It is the suppression of the activity of PNP in the second experimental groups of all organs.

**Fig. 1.**
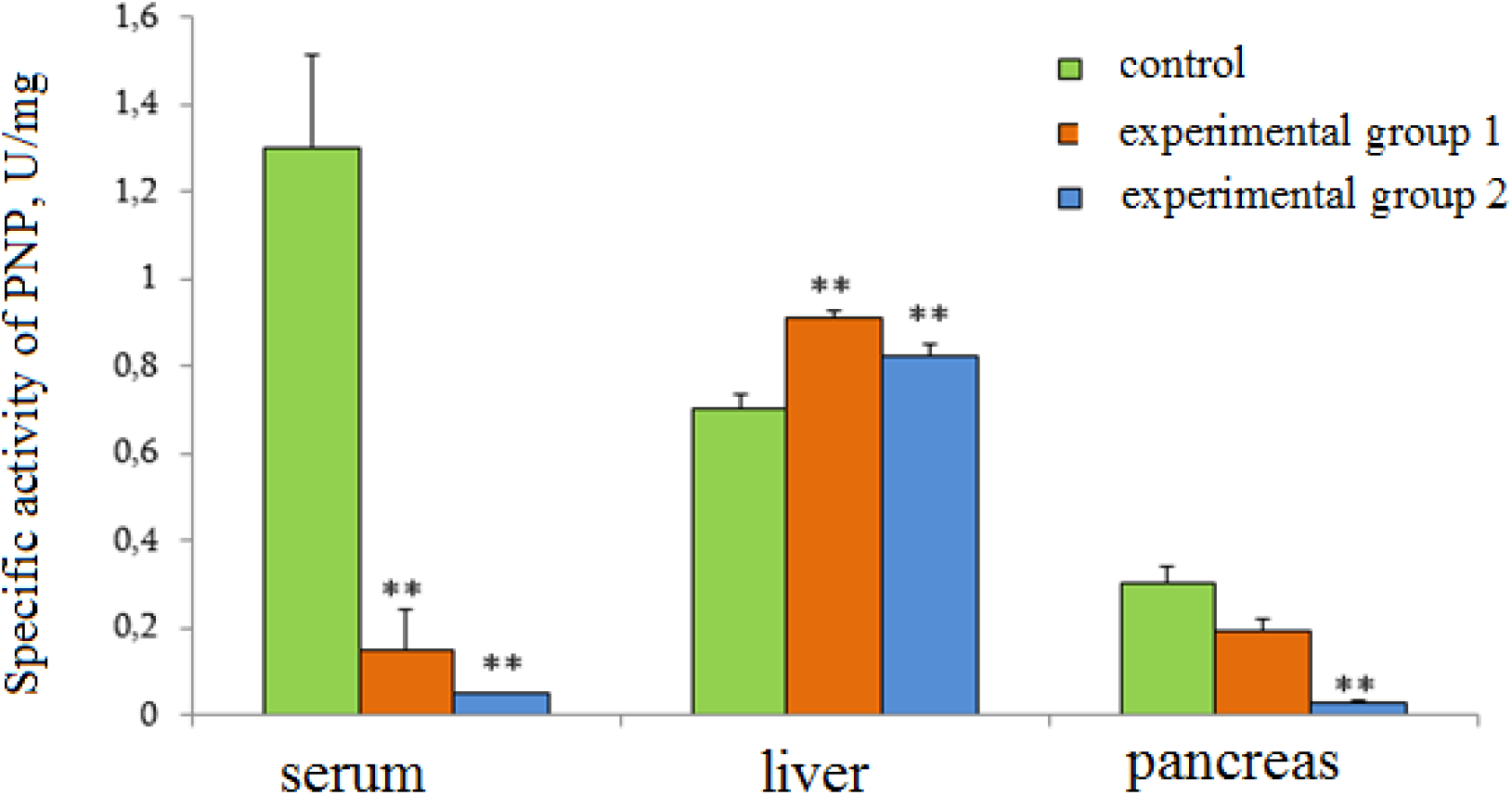
Dynamics of changes in the activity level of PNP in blood serum, liver, pancreas in both experimental groups: experimental groups 1-10 days after alloxan injection, experimental group 2-20 days after alloxan injection. *Note*\*\* - Difference from control is statistically significant at *p* ≤ 0.01

In our previous works we investigated the influence of different factors of the environment on PNP activity in some organs of rats. Different degree of sensitivity of the investigated organs, assessing the meanings of enzymatic activity on changeable environmental conditions. Blood serum was the most affected, and hepatic PNP showed resistant properties [8].

In this work the activity of PNP in all organs changed/ was changing in different ways. On the diagram we can see a dramatic decrease in the level of enzymatic activity in the blood serum of the first experimental group, compared to the control and almost complete suppression of activity in the 2^nd^ one (Fig. 1). Such a persistent / steady decline in the PNP activity is characteristic for a range of pathologies with significant / expressed immunodeficiency [1]. Moreover one can not exclude / it can not be excluded, that decline in the activity of serum enzyme can be connected with the disruption of blood cells (especially erythrocytes), which have high PNP activity [4].

In contrast to blood serum, the meanings of PNP activity levels in the liver of both experimental groups rise. And although the activity in the second experimental group is somewhat reduced compared to the first one, nevertheless it remains elevated compared to the control group of animals. It is well-known that liver plays a vital role in the synthesis and metabolism of carbohydrates, lipids and proteins, hence it has a very wide functional and metabolic spectrum. The organ also consists of hepatocytes, where the activity of PNP is the highest. Therefore, when there is a disruption in the glucose metabolism and synthesis in diabetes mellitus, we can only assume that PNP activity level will increase.

Pancreas is a unique organ, which has both exocrine and endocrine function. There are different methods for revealing the impairment of the exocrine function of pancreas, including assessment of enzymatic activity or assessment of the cleavage degree of the substrate by pancreatic enzymes [7]. The role of immunological research for assessment of the severity of secondary immunodeficiency in various diseases of the pancreas has increased. The results of studies of immune parameters in such patients indicated different deviations in the immune system, which means that the mechanisms of natural immune resistance were weakened. Hyperglycemia, as a consequence of diabetes mellitus, causes severe damage to various tissues and hence changes in the activity of enzymes in these tissues [9].

The results of our research about revealing of differences in PNP activity in pancreas on the model of alloxan diabetic rats show that at the later stages of the disease (in our experiment it is 20 days after alloxan injection) the activity of PNP of second experimental group is completely suppressed, which indicates about significant changes in the immune status of the organism.

As the results of our studies showed, activation of the processes of the free radical oxidation (MDA), decrease of the activity of glutathione peroxidase and also increase of nitric oxide in blood plasma are noted /observable in the pancreatic tissue of alloxan diabetic rat /in the tissue of pancreas of alloxan diabetic rats on the 20^th^ day of diabetes development (Fig .2)

**Fig. 2.**
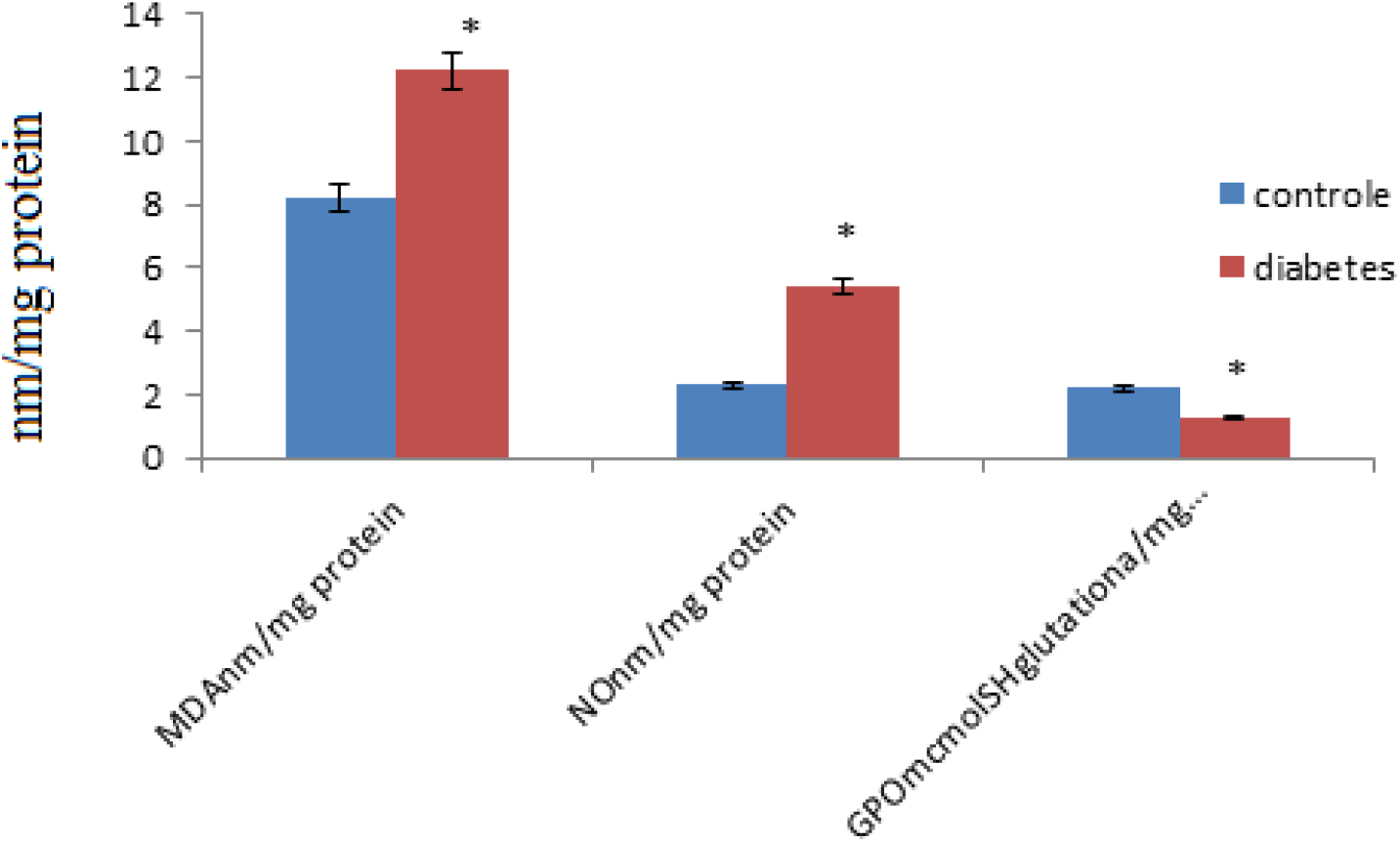
Nitric oxide content in blood plasma (μmol/g of protein), gluthatione peroxidase activity (μmol SH of gluthatione / mg of protein) and products of lipid peroxidation MDA (nmol/g of protein), in the mitochondrial fraction of pancreas in normal state and at alloxan diabet (n=15) Note: * - the difference from the control is significant when P<0.05

The increase in the content of lipid peroxides in the mitochondrial fraction of the pancreas of alloxandiabetic rats is facilitated by the high content of easily oxidizable substrates in it, such as polyunsaturated fatty acids, as well as the presence of non-heme iron in the cytochromes of the respiratory chain, which are activators of LPO.

In these conditions the mitochondrial electron transfer chain becomes a powerful source of formation of reactive oxygen species – non-stable and extremely reactive metabolites. The disruption of normal functioning of the mitochondrial electron transfer chain is also facilitated by activation of nitric oxide formation. The research on nitric oxide content revealed elevation of its content in the mitochondrial fraction of animals with alloxan diabetes (Fig. 2). When nitric oxide (NO) interacts with the superoxide anion, the active compounds nitrosonium (NO+), nitroxyl (NO−) and peroxynitrite (ONOO) are formed. At high concentrations NO (>1 mkm) interacts with the complexes of the respiratory chain (cytochrome oxidase, ubiquinone) leading to the suppression of the synthesis of adenosine triphosphate [13]. Targets of direct action of NO are copper and zinc atoms, which are parts of the enzymes. It is estimated that nitric oxide, which forms in the islets and beta cells of pancreas, plays an important role in the mechanisms of disruption and death of beta cells, which leads to a dramatic decrease in their amount and DM development [14].

The processes of free radical oxidation in the organism are controlled by the antioxidant system. Easily oxidizing peptides play an important role in the antioxidant protection of the organism, glutathione - a thiol containing tripeptide, formed by amino acids cysteine, glutamic acid and glycine, has a special role among these compounds.

Glutathione in complex with glutathione peroxidase and glutathione reductase makes a system which has an important role in the regulation of metabolic processes in cells, the main function of these enzymes is the reduction of hydroperoxides to alcohols [15].

As the results of the research show (Fig. 2), there is a decrease in the activity of glutathione peroxidase in alloxan diabetes.

## CONCLUSIONS

The obtained results have demonstrated statistically significant changes in the PNP activity level indicators in the investigated organs and stages of the disease. The indicators of oxidative processes MDA, NO and changes of the activity of GPx and PNP can be used as additional paraclinical indicators for early diagnostics and for detection and refinement of the level of activity of the pathologic process, its development and stage of the disease, which contributes to the appointment of individualized adequate therapy.

## References

1. Eyal Grunebaum, Nicholas Campbell, Matilde Leon-Ponte, Xiaobai Xu and Hygo Chapdelaine. Partial purine nucleoside phosphorylase deficiency helps determine minimal activity required for immune and neurological development. Front. Immunol., 2020: 11: 1257. doi:10.3389/fimmu.2020.01257.

2. Linden J., Koch-Nolte F., Dahl G. Purine release, metabolism and signaling in the Inflammatory response. Annu. Rev. Immunol., 2019, V.37, P. 325–347. doi:10.1146/annurev-immunol-051116-052406.

3. Panevin T.S., Zhelyabina O.V., Eliseev M.S., Shestakova M.V. Urate-lowering effects dipeptidyl pepridase-4 inhibitors. Diabetes Mellitus, 2020: V.23, N4, P. 349–356. doi:10.14341/DM12412.

4. Bzowska A., Kulikovcka E., Shugar D. Purine nucleoside phosphorylase: properties, functions and clinical aspect// Pharmacol. Ther. 2000, V.88, N3, P.349–425 doi:10.1016/50163-7258(00)00097-8.

5. Amacher D.E., Adler R., Herath A. et al. Use proteomic methods to identify or hypertrophy// Clin.Chem. 2005, V.51, N10, P.1796–1803. doi:10.1373/clinchem.2005.049908

6. Pogosian L.H., Akopian J.I. Purine nucleoside phosphorylase. Biomed. Chem. 2013, V.59, N5, P.483–497. doi:10.18097/pbmc20135905483.

7. Piciucchi M., Capurso G., Archibugi L., Delle Fave MM., Capasso M., Delle FG. Exocrine pancreatic insufficiency in diabetic patients: prevalence, mechanisms and treatment. Int. J.Endocrinol. 2015; 2015:595649. doi:10.1155/2015/595649.

8. Poghosyan L.H., Akopyan J.I. et al. Influence of electromagnetic radiation with frequency of 900 and 1800 MHz on the activity of purine nucleoside phosphorylase and Alkaline phosphatase in some organs of the Rats. Medical radiology and radiation safety. 2016, V.61, N 6, P.5–10.

9. Dominquez-Minoz J.E. Chronic pancreatitus and persistent steatorrhea:what is the correct dose of enzymes? Clin.Gactroenterol.Hepatol., 2011; V.9, N7, P.541–546. doi:10.1016/j.cgh.2011.02.027.

10. K Luc 1, A Schramm-Luc 1, T J Guzik 1 2, T P Mikolajczyk 1 3. Oxidative stress and inflammatory markers in prediabetes and diabetes. J Physiol Pharmacol. 2019 Dec;70(6). doi: 10.26402/jpp.2019.6.01. Epub 2020 Feb 19. DOI: 10.26402/jpp.2019.6.01

11. O. Macdonald I. Molecular pathways associated with oxidative stress in diabetes mellitus. Biomed Pharmacother. 2018 Dec;108:656–662. doi: 10.1016/j.biopha.2018.09.058. Epub 2018 Sep 20. DOI: 10.1016/j.biopha.2018.09.058

12. A.V. Razygraev. Method of Determination of Glutathione Peroxidase Activity Kliniko-Laboratornyi Konsilium. 2004. – № 4. – C. 19–22.

13. Guy C Brown, Vilmante Borutaite.Nitric oxide inhibition of mitochondrial respiration and its role in cell death. Free Radic Biol Med. 2002 Dec 1;33(11):1440–50. doi: 10.1016/s0891-5849(02)01112-7.

14. J. Pfeilschifter, Does nitric oxide, an inflamatory mediator of glomerular mesangial cells, have a role in diabetic nephropathy? Kidney Int.,48 (51), 50 – 61 (1995)

15. A V Razygraev, A D Yushina, I A Titovich. A Method of Measuring Glutathione Peroxidase Activity in Murine Brain in Pharmacological Experiments. Bull Exp Biol Med. 2018 Jun;165(2):292–295. doi: 10.1007/s10517-018-4151-5. Epub 2018 Jun 20.

16. J. Sochor, B. Ruttkay-Nedecky, +3 authors R. Kizek. Automation of Methods for Determination of Lipid Peroxidation Published Chemistry 29 August 2012. DOI:10.5772/45945 Corpus ID: 871574

17. M.V Mazhitova, SPECTROPHOTOMETRIC DETERMINATION OF NITROGEN MONOXIDE METABOLITE LEVEL IN BLOOD AND BRAIN PLASMA IN WHITE RATS // Modern problems of science and education. – 2011. – № 3.; URL: https://science-education.ru/ru/article/

